# Metabotropic glutamate group II receptor activation in the dorsal striatum suppresses the motivation for cocaine in rats

**DOI:** 10.1101/2022.09.16.507381

**Authors:** Shaun Yon-Seng Khoo, Anne-Noël Samaha

## Abstract

After a history of intermittent cocaine intake, rats develop patterns of drug use characteristic of addiction. The dorsal striatum is involved in the increased pursuit of cocaine after intermittent drug self-administration experience. Within the dorsal striatum, chronic cocaine use changes metabotropic glutamate type II receptor (mGlu_2/3_) density and function. We examined the extent to which activity at these receptors mediates responding for cocaine after intermittent cocaine use. In Experiment 1, male Wistar rats (n = 11) self-administered 0.25 mg/kg/infusion cocaine during 10 daily intermittent access (IntA) sessions (5 min ON/25 min OFF, for 5 h/session). We then examined the effects of intra-dorsal striatum infusions of the mGlu_2/3_ receptor agonist LY379268 (0, 1, and 3 μg/hemisphere) on cocaine self-administration under a progressive ratio schedule of reinforcement. We observed a non-significant tendency for LY379268 to reduce responding for cocaine. In Experiment 2, we used a larger sample of male (n = 11) and female (n = 10) rats. Across 10 IntA sessions, the sexes showed similar levels of cocaine intake. Across the sexes, locomotion significantly increased over sessions, suggesting that rats developed psychomotor sensitization to self-administered cocaine. After 10 IntA sessions, intra-dorsal striatum LY379268 significantly reduced breakpoints achieved for cocaine, active lever presses, and cocaine infusions earned under progressive ratio. LY379268 had no effects on locomotion or inactive lever presses, indicating no motor effects. These results suggest that mGlu_2/3_ receptor activation in the dorsal striatum suppresses incentive motivation for cocaine, and this holds promise for anti-addiction therapeutics.

## Introduction

Drug addiction (or substance use disorder) is amongst the leading preventable causes of death and illness across the world. Regular or problematic cocaine use is associated with significantly elevated mortality and Canada and the United States have among the highest prevalence of cocaine use disorder in the world (Peacock et al., 2018). Psychosocial interventions, such as cognitive behavioural therapy and contingency management, have some efficacy in treating drug addiction (Minozzi et al., 2016), and it has been suggested that best practices should include combined pharmacological and behavioural interventions (Anton et al., 2006; Kampman & Jarvis, 2015; Carroll, 1997; Ray et al., 2020). However, there are still no approved pharmacotherapies for cocaine use disorder (Brandt et al., 2021). It is therefore imperative to study the brain mechanisms involved in addiction-relevant patterns of cocaine use to better understand the neural mediators of the addiction process and identify potential therapeutic targets.

To study brain mechanisms involved in addiction, our laboratory employs an intermittent access (IntA) cocaine self-administration procedure recently developed in rats (Zimmer et al., 2012). This more closely models human patterns of cocaine use, which tend to be intermittent both between and within bouts of drug use (Zimmer et al., 2012; Samaha et al., 2021; Kawa et al., 2016, 2019; Allain et al., 2015; Cohen & Sas, 1994; Leri et al., 2004; Beveridge et al., 2012). Compared to self-administration procedures where cocaine is available continuously (Short Access or Long Access procedures), IntA to cocaine is also uniquely effective in producing addiction-like behaviours in rats. This includes a persistent increase in incentive motivation to obtain cocaine (Calipari et al., 2014; Allain & Samaha, 2019; James et al., 2019; Algallal et al., 2020), enhanced drug taking despite aversive consequences (James et al., 2019; Singer et al., 2018), and especially robust cue-induced relapse-like behaviour (Kawa et al., 2016, 2019; James et al., 2019; Singer et al., 2018; Gueye et al., 2018). Therefore, findings from studies using IntA self-administration may, theoretically, be readily translatable into new clinical approaches.

Several previous studies have implicated the dorsal striatum in incentive motivation for cocaine after a history of IntA cocaine use. IntA cocaine-taking experience increases cocaine-induced gene regulation in the dorsal striatum (Minogianis & Samaha, 2020). This is not just a correlational effect, because after IntA cocaine self-administration, temporary, pharmacological inhibition of the dorsal striatum with baclofen and muscimol microinjection reduces responding for cocaine in progressive ratio (PR) tests (Minogianis et al., 2019). Chemogenetic inhibition of dorsal striatum medium spiny neurons projecting to the substantia nigra (i.e., direct pathway neurons) also suppresses cue-induced cocaine seeking, specifically in IntA rats that have developed a robust addiction phenotype (Yager et al., 2019). These findings are consistent with previous work using Short Access and Long Access cocaine self-administration sessions, where intra-dorsal striatum baclofen and muscimol suppressed cocaine-seeking after abstinence (Pacchioni et al., 2011). The dorsal striatum has therefore been implicated in motivation for cocaine by multiple studies using IntA and other self-administration models.

Within the dorsal striatum, chronic cocaine use changes metabotropic glutamate type II receptor (mGlu_2/3_) density and function. mGlu_2/3_ receptors are primarily presynaptic, G_*i*/*o*_-coupled receptors and they typically function to suppress synaptic glutamate signalling (Conn & Pin, 1997; Imre, 2007; Schoepp, 2001). Electrophysiological and pharmacological studies show that in the striatum and other brain regions, mGlu_2/3_ receptor agonists reduce glutamate release and excitatory postsynaptic potentials (Kilbride et al., 1998; Dohovics et al., 2003). In both rats (Pomierny-Chamiolo et al., 2017) and non-human primates (Beveridge et al., 2011), cocaine self-administration significantly increases mGlu_2/3_ receptor expression and functional activity in the dorsal striatum. This is consistent with the effects of chronic cocaine self-administration on mGlu_2/3_ receptors in several other cortical and limbic-related brain regions (Allain et al., 2017; Hao et al., 2010). mGlu_2/3_ receptor agonists, given systemically (Allain et al., 2017; Hao et al., 2010; Baptista et al., 2004; Adewale et al., 2006; Karkhanis et al., 2016) or into brain nuclei such as the nucleus accumbens (Peters & Kalivas, 2006) can reduce cocaine-seeking and -taking behaviours. To our knowledge, however, no published study has examined the effects of activating mGlu_2/3_ receptors in the dorsal striatum on cocaine use.

Thus, the aim of the present study was to determine how activating mGlu_2/3_ receptors in the dorsal striatum with a selective agonist (LY379268) influences incentive motivation for cocaine, as measured by responding for the drug under a progressive ratio schedule of reinforcement. We assessed this in rats with an IntA cocaine self-administration history, to model the intermittent patterns of cocaine use reported in humans (Zimmer et al., 2012; Samaha et al., 2021; Kawa et al., 2016, 2019; Allain et al., 2015; Cohen & Sas, 1994; Leri et al., 2004; Beveridge et al., 2012). Based on the available literature, we hypothesised that microinjections of LY379268 into the dorsal striatum will decrease progressive ratio responding for cocaine.

## Methods

### Animals

Adult Wistar rats were obtained from Charles River Laboratories. For experiment 1, 20 males weighing 225-250 g on arrival were received from Charles River Canada (Area C62, Saint-Constant, QC). Due to supplier restructuring, experiment 2 used 17 males weighing 225-250 g and 17 females weighing 150-175 g on arrival from Charles River Raleigh (Area R06, NC, United States). Rats were housed in clear polycarbonate cages (44.5 cm × 25.8 cm × 21.7 cm) in a climate-controlled colony room (22 ± 1°C, 30 ± 10% humidity). They were housed individually to prevent damage to their catheter and intracerebral implants and were maintained on a reverse 12 h:12 h light/dark cycle (lights off at 8:30 am). Rats were trained and tested during the dark phase of their cycle. After arrival, rats were handled for at least 3 days before the commencement of behavioural procedures. Rats had unlimited water in their home-cage. They had unlimited chow (Rodent 5075, 18% protein, 4.5% fat, Charles River Laboratories) until 2 days before the commencement of food self-administration training, at which point they were restricted to 25 g/day for males and 20 g/day for females. This feeding schedule corresponds to 70-80% of what ad libitum-fed rats would eat (Algallal et al., 2020). Such mild food restriction produces healthier rats compared to ad libitum feeding, which promotes excessive fat deposition and obesity (Martin et al., 2010; Rowland, 2007). All experiments were approved by the animal ethics committee at the Université de Montréal and conducted in accordance with guidelines from the Canadian Council on Animal Care.

### Behavioural Apparatus

Behavioural training took place in 30 identical standard operant chambers (Med Associates Inc, St-Albans, VT, United States) with standard grid floors. The left-side wall was equipped with a 100 mA white incandescent house light and a 2,900 Hz Sonalert tone generator (ENV-223AM) calibrated to 85 dB. The right-side wall was equipped with a food cup flanked on either side by a retractable lever (ENV-112CM) calibrated to minimum tension for maximum sensitivity. A white cue light (ENV-221M, 100 mA incandescent bulb) was located above each lever. During food self-administration training, 45-mg grain pellets (F0165, BioServ, Flemington, NJ, United States) were delivered to the food cup via a pellet dispenser (ENV-203-45). To enable cocaine self-administration, a spring tether was fed through the top of the chamber, connected via Tygon® tubing (0.02” ID, 0.06” OD) to a 20-mL syringe placed on a syringe pump (PHM-100, 3.33 RPM).

### Drugs

Cocaine hydrochloride (Galenova, St-Hyacinthe, QC, Canada) was dissolved in 0.9% saline and vacuum-filtered. Cocaine concentrations were adjusted every 3-4 days as required according to the average bodyweight of the animals, with separate solutions for males and females. The mGlu_2/3_ agonist LY379268 was obtained from Tocris (Cat#: 2453, Batch#: 9B/232416 & 10A/254948, CAS: 191471-52-0, Bio-Techne, Toronto, ON, Canada) and dissolved in an artificial cerebrospinal fluid vehicle (for recipe see “aCSF (1x)”, 2011) by gentle heating in a water bath and sonication.

### Food Self-Administration Training

Figure 1a presents an outline of behavioural training and testing. Two days prior to food self-administration training, rats were restricted to 25 g/day for males and 20 g/day for females. Rats were then allowed to lever press for food pellets during daily sessions. Responses on the lever to the left of the food cup (active lever) were reinforced with a food pellet on a fixed ratio-1 (FR1) schedule. Responses on the right lever (inactive lever) had no programmed consequences. Each pellet delivery was accompanied by retraction of both levers, 5 s of illumination of the cue light above the active lever and 5 s of tone presentation. After presentation of the 5-s tone-light cue, the levers remained retracted for a further 20-s time-out period before extending again. Each session ended after rats earned 100 pellets or 1 h had elapsed.

**Figure 1.**
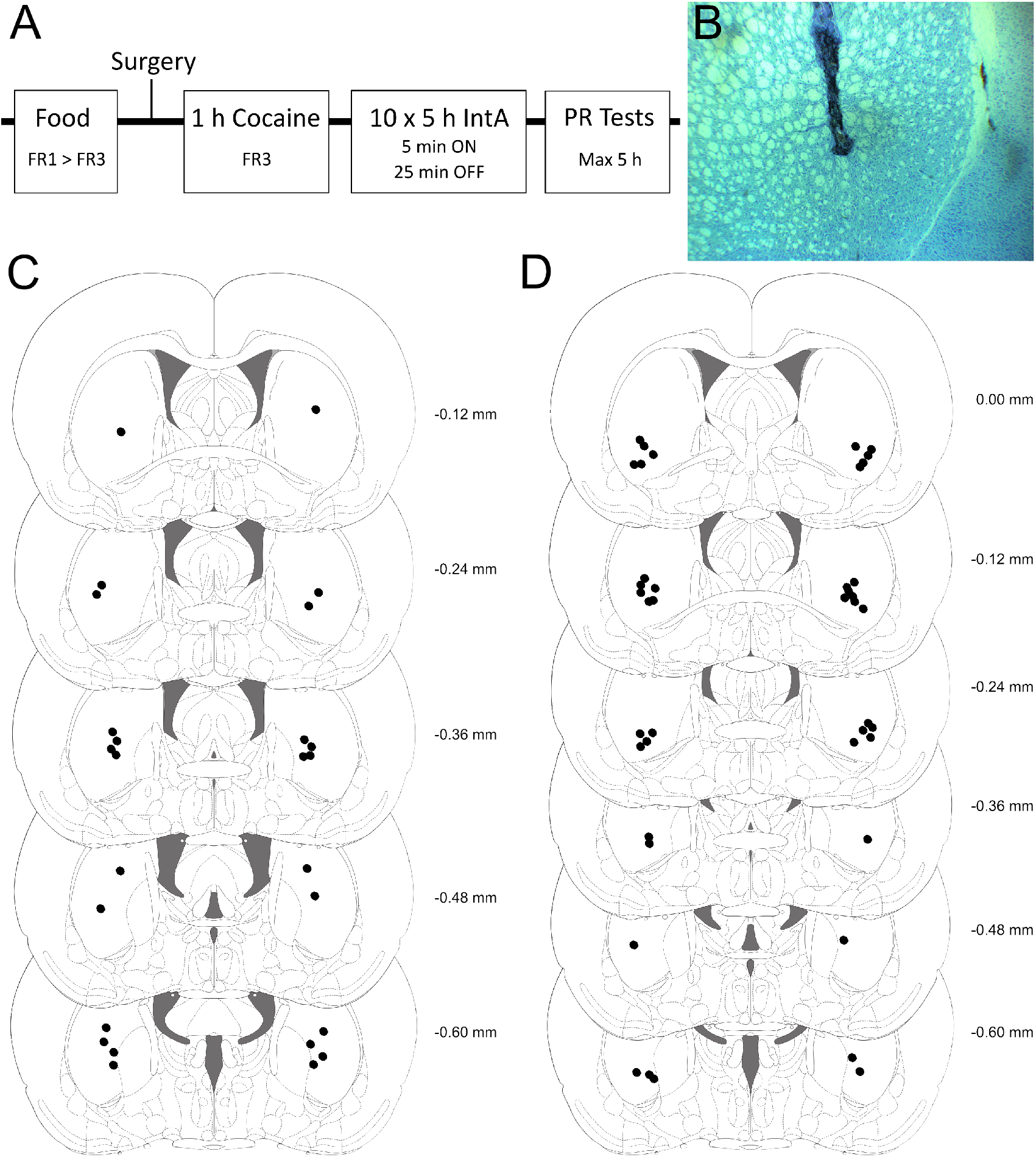
Experimental design and histology. (a) Experiments involved first providing food self-administration training under fixed ratio-1 (FR1) then fixed ratio-3 (FR3) schedules. This was followed by surgical implantation of jugular vein catheters and bilateral cannulae into the dorsal striatum. After recovery, rats received 1-h cocaine self-administration training sessions, and then 5-h intermittent access (IntA) cocaine self-administration. Finally, rats could respond for cocaine under a progressive ratio schedule of reinforcement (PR) following bilateral vehicle and LY379268 microinjections into the dorsal striatum. PR tests lasted a maximum of 5 h. (b) A representative photomicrograph of a thionin-stained brain section showing microinjector placement in the dorsal striatum. Estimated microinjection placements for (c) experiment 1 and (d) for experiment 2 illustrated on brain sections from Paxinos & Watson (2007).

When rats earned at least 20 pellets during an FR1 session, they progressed to FR3 sessions for days 3-5. Rats that did not meet the 20-pellet criterion were given an overnight session with no maximum number of pellets. In experiment 1, all rats met the acquisition criterion in a maximum of two FR1 sessions, and none required overnight sessions. In experiment 2, after two FR1 sessions, 6 male and 6 female rats required overnight sessions, and 2 males and 3 females were excluded from the experiment for not meeting the 20-pellet criterion after 5 days of food self-administration training at FR3.

### Surgery

To enable cocaine self-administration and intracerebral microinjections, respectively, jugular vein catheters were implanted as previously described (Samaha et al., 2011) and cannulae were implanted bilaterally into the dorsal striatum (Minogianis et al., 2019). Rats received both implants during a single surgery. Rats were first deeply anaesthetised using isoflurane, sites of interest were shaved, and rats were given 22,000 IU/kg i.m. penicillin (CDMV, St-Hyacinthe, QC, Canada) and 5 mg/kg s.c. carprofen (Rimadyl, CDMV). Catheters were implanted in the left jugular vein with a back port exiting between the shoulder blades. Catheters were flushed with 0.05 – 0.1 mL of a 0.2 mg/mL heparin and 2 mg/mL enroflaxin (Heparin: Sigma-Aldrich, Oakville, ON, Canada; Enroflaxin: CDMV) solution. Catheters were then sealed with a dummy and rats were moved to a stereotaxic frame. Twenty-two gauge cannula (P1Technologies, Roanoke, VA, United States) were then implanted bilaterally into the dorsal striatum with coordinates based on Minogianis et al. (2019): from bregma, A/P – 1.2, M/L +/-3.7 and D/V – 5.6 mm. Cannulae were secured to the skull using four screws and dental acrylic before being sealed with obturators that were flush with cannulae tips. Injectors would later extend 2 mm beyond cannulae tips, to D/V -7.6 mm. Wounds were swabbed with chlorhexidine and Flamazine® cream (1% w/w silver sulfadiazine) after surgery and for the first 3 days of recovery (and then as required). Post-surgery catheter maintenance involved flushing on alternate days with 0.05 – 0.1 mL saline or heparine/enroflaxin and continued until the end of the experiment. Rats were given at least one week to recover from surgery before cocaine self-administration training.

### Cocaine Self-Administration Training

Rats received 6-10 days of 1-h cocaine self-administration training sessions. These sessions functioned identically to food self-administration sessions, except that rats now received cocaine (0.25 mg/kg/infusion in approx. 0.14 mL) instead of food pellets. Cocaine was delivered on an FR3 schedule of reinforcement. Each cocaine injection was accompanied by lever retraction, a 5-s light+tone cue, and an additional 20-s timeout period, after which levers were again inserted into the cage. Rats needed to earn at least 6 infusions, evenly spaced over the hour, after session 6 to pass to the intermittent-access cocaine self-administration sessions described next. In experiment 1, all rats met the acquisition criteria in 6 sessions. In experiment 2, 6 male and 6 female rats were given between 1 and 4 additional sessions and 2 male rats were then excluded for not meeting criteria after 10 sessions.

### Intermittent Access Cocaine Self-Administration

Rats then received 10 sessions of intermittent access to cocaine (0.25 mg/kg/infusion). As previously described (Zimmer et al., 2012; Algallal et al., 2020; Allain et al., 2017, 2018), intermittent access (IntA) sessions were 5-h long, with 12 cycles of 30 min. Each cycle included a 5-min cocaine ON period during which both the active and inactive levers were presented, and cocaine was available on an FR3 schedule of reinforcement, with a 20-s timeout period following each self-administered injection. Each cocaine injection was paired with a 5-s tone+light cue, and levers were retracted during each 5-s infusion and 20-s timeout period (i.e., for a total of 25 s). At the end of each 5-min cocaine period, levers were retracted (or remained retracted, if during a timeout period) for a 25-min cocaine OFF period, until the start of the next cycle.

### Habituation to Microinjection Procedures

At the conclusion of the 10 IntA self-administration sessions, rats were kept in their home cages for 3-5 days. During this time they were habituated to microinjection procedures in the colony room. Briefly, dummies were removed and replaced and rats were minimally restrained to prevent grooming of the cannula for a gradually increasing duration up to 5 min to simulate microinjection and infusate diffusion time.

### Catheter Patency

The day after the last IntA session, and after the progressive ratio testing described next, catheters were tested for patency by injecting 0.05 mL of propofol (10 mg/mL, i.v.; Fresenius Kabi Canada Ltd, Richmond Hill, ON, Canada). Ataxia within 5-10 s of propofol administration indicated functional catheters. Rats that did not become ataxic during this time frame were re-tested immediately, and, if they failed again they were excluded from further behavioural testing.

### Responding for Cocaine under a Progressive Ratio Schedule of Reinforcement

To measure the motivation to obtain cocaine, we quantified breakpoints achieved for cocaine under a progressive ratio schedule of reinforcement (PR). The rats were presented with the active and inactive levers and cocaine injections were available under an exponentially escalating ratio according to the following formula (round[5e^(*reinforcer number* × 0.2)^ – 5]), devised by Richardson & Roberts (1996). Each earned cocaine injection was paired with a 5-s tone+light cue, and levers were retracted during each 5-s infusion and 20-s timeout period (i.e., for a total of 25 s). Session ended either 1 h after the last self-administered cocaine injection or after 5 h had elapsed. Breakpoint was defined as the last successfully completed ratio. Previous microinjection studies in rats show that LY379268 significantly reduces drug-seeking behaviour across 3-h test sessions, suggesting that the compound is effective for at least 3 h (Bossert et al., 2006). Thus, we used 0.125 mg/kg/infusion cocaine for our PR tests here, because rats typically reach break-point at this dose in around 3 h (Minogianis & Samaha, 2020). Rats received at least one day off between PR test sessions.

### Experiment 1

Rats received aCSF, 1, and 3 μg/side of LY379268 using a within-subjects latin square design (1 treatment/PR test session). Prior to each PR test session, the experimenter minimally restrained the rats and microinjected 2 μL aCSF or LY379268 into the dorsal striatum, at a rate of 0.5 μL/min using a syringe pump (Harvard Apparatus, Saint-Laurent, QC, Canada). Injectors were left in place for an additional minute (i.e. 5 min total). PR testing began immediately after microinjections. On the second PR test session, during which some rats had received aCSF, some 1 μg and some 3 μg/side LY379268, breakpoints were significantly lower than we typically observe at 0.125 mg/kg/infusion cocaine. This was observed across all 3 microinjection types (aCSF, 1 and 3 μg/side LY379268). For this reason, the next day we gave all rats a baseline PR session without microinjections, and the second test was then repeated. Thus, all rats received a total of 4 microinjections.

### Experiment 2

All rats received 5 PR test sessions and 4 microinjections/hemisphere. On the first session, all rats received 2 μL aCSF immediately prior to PR testing. On the 2nd session, the rats were tested under baseline PR conditions, with no microinjections. This allowed us to determine whether aCSF microinjection to previously microinjection-naïve rats influences responding for cocaine. During sessions 3-5, rats were tested with aCSF, 1 μg, and 3 μg/side LY379268, under a within-subjects latin square design.

### Histology

Within one week of the completion of testing, rats were anaesthetised and decapitated. Brains were dissected and frozen at - 20°C in a cryostat (Leica CM1850, Leica Biosystems, IL, United States). They were then sectioned coronally at 40 μm and thaw-mounted on glass microscope slides. Sections were then stained using thionin. Injector placement was verified while blind to experimental results and placements were mapped according to the rat brain atlas (Paxinos & Watson, 2007). Figure 1b shows a representative photomicrograph at 4x magnification, and Figures 1c and 1d show estimated injector placements for experiments 1 and 2, respectively. No rats were excluded following histology.

### Data Analysis

Data were analysed using SPSS 27 (IBM, Armonk, NY, United States). Repeated measures ANOVAs were used with Bonferroni-corrected post-hoc comparisons following significant effects. We observed dose order effects in Experiment 2, where receiving LY379268 in one test session suppressed responding in subsequent tests. For this reason, we compared data from the first test with a given dose of LY379268 (i.e., in rats that were LY379268-naïve before this test) to the very first aCSF test, administered before any LY379268 testing. For this, results were therefore analysed as a mixed-design ANOVA with vehicle vs LY379268 as a within-subjects factor (‘Treatment’) and dose of LY379268 as a between-subjects factor (‘Dose’). The Greenhouse-Geisser correction was applied to degrees of freedom when ε < 0.75.

## Results

### Experiment 1

***Cocaine self-administration during intermittent access sessions*** Rats (n = 11) that had acquired reliable food and then cocaine self-administration during training now received ten, 5-h IntA cocaine self-administration sessions. As shown in Figure 2a, rats responded for cocaine at high levels throughout the 10 IntA sessions, and lever pressing remained stable across sessions (main effect of Session; F(3.12,31.16) = 0.68, *p* = 0.58, ε = 0.35). The rats discriminated between levers, with a marked preference for the active lever (main effect of Lever; F(1,10) = 15.86, *p* = 0.003). Lever discrimination also remained stable across IntA sessions (no significant Lever × Session interaction effect; F(2.42,24.18) = 1.11, *p* = 0.36, ε = 0.27). Accordingly, the number of self-administered cocaine infusions was also stable across IntA sessions (Main effect of Session; F(9,90) = 0.98, *p* = 0.46; Figure 2b). This steady consumption resulted in linear cumulative cocaine intake (Figure 2c). Thus, rats reliably self-administered cocaine under IntA conditions, and drug consumption remained stable across sessions.

**Figure 2.**
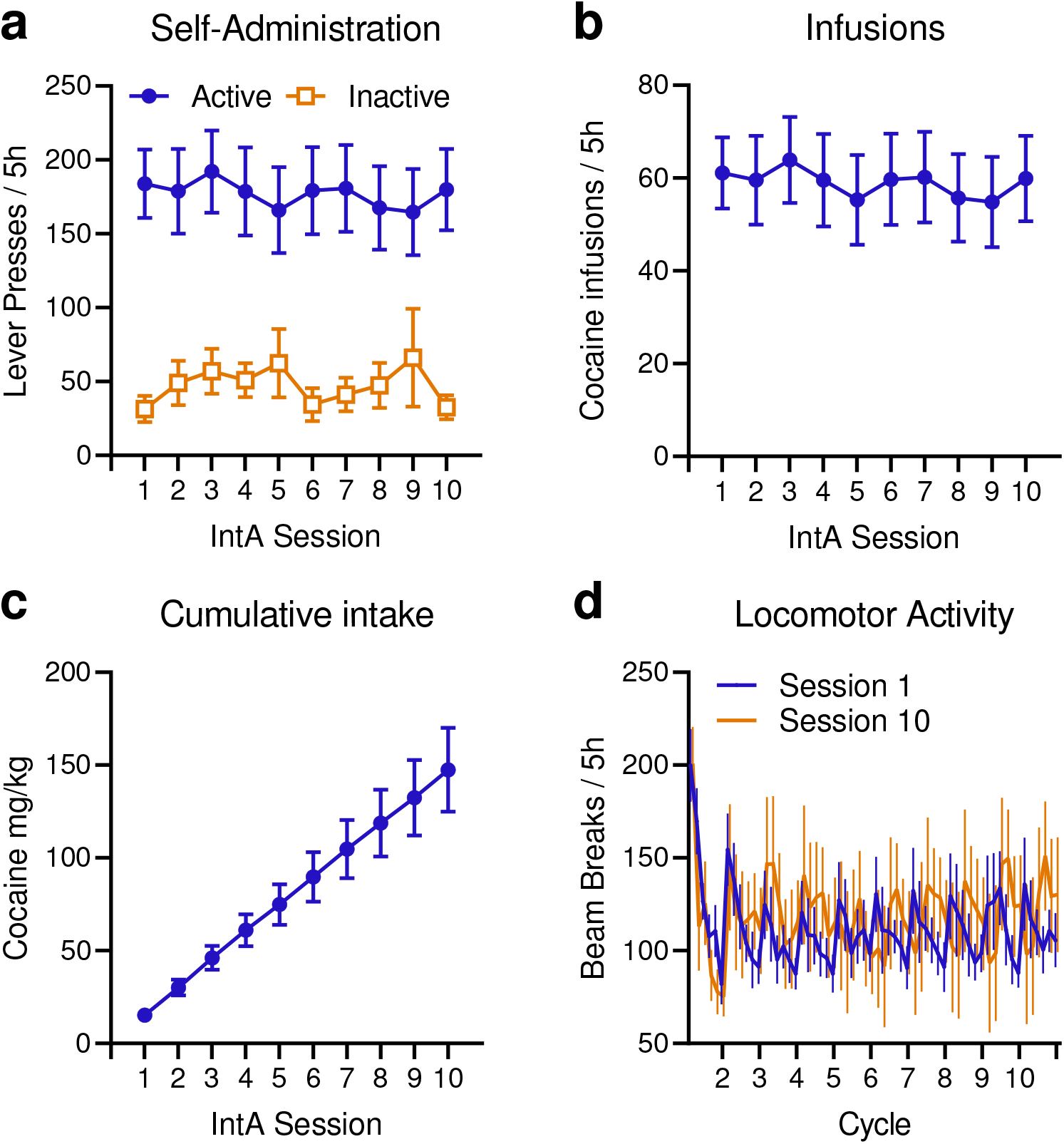
Intermittent access cocaine self-administration during Experiment 1. (a) Rats (n = 11 males) showed high and stable levels of instrumental responding for cocaine, with clear discrimination between active and inactive levers. (b) The rats earned a stable number of cocaine infusions across IntA sessions, resulting in (c) a linear increase in cumulative cocaine intake across self-administration sessions. (d) During the 1st and 10th IntA sessions, locomotor behaviour followed a spiking pattern, peaking at the beginning of each cocaine ON phase and decreasing during each cocaine OFF phase. Data are shown as means ± SEM.

Figure 2d shows the number of locomotor beam breaks for IntA session 1 and IntA session 10. At the beginning of each 5-min cocaine ON period, locomotor activity rapidly rises, before gradually decreasing during the 25-min cocaine OFF periods. Thus, spikes in locomotor activity coincide with cocaine availability during IntA sessions. There were no statistically significant changes in locomotion from the first to the last IntA session. Indeed, paired t-tests revealed no significant difference in average locomotor activity peaks (in beam breaks / 5 min bin) between session 1 (Mean = 138.78, SEM = 16.72) and session 10 (Mean = 146.82, SEM = 25.37; t(10) = 0.35, *p* = 0.73).

#### Effects of LY379268 on responding for cocaine under a progressive ratio schedule of reinforcement

Starting 3-5 days after the last IntA session, we assessed the effects of bilateral intra-dorsal striatum infusions of LY379268 on cocaine self-administration under a PR schedule of reinforcement. There were nonsignificant trends towards a dose-related decrease in breakpoints (i.e., last ratio reached; (F(2,20) = 2.96, *p* = 0.075; Figure 3a) and total number of active lever presses during the PR test session (F(2,20) = 3.3, *p* = 0.058; Figure 3b). There was no significant effect of Treatment on the number of cocaine infusions earned (F(2,20) = 1.83, *p* = 0.19; Figure 3d). Importantly, LY379268 had no significant effect on inactive lever presses (F(2,20) = 0.73, *p* = 0.49; Figure 3c) or on locomotor activity as measured by beam breaks per min (F(2,20) = 1.93, *p* = 0.17; Figure 3e). This suggests that the compound did not suppress motor behaviour non-specifically. Finally, as shown in Figure 3f, there was also no effect of Treatment on session length (F(2,20) = 0.71, *p* = 0.5).

**Figure 3.**
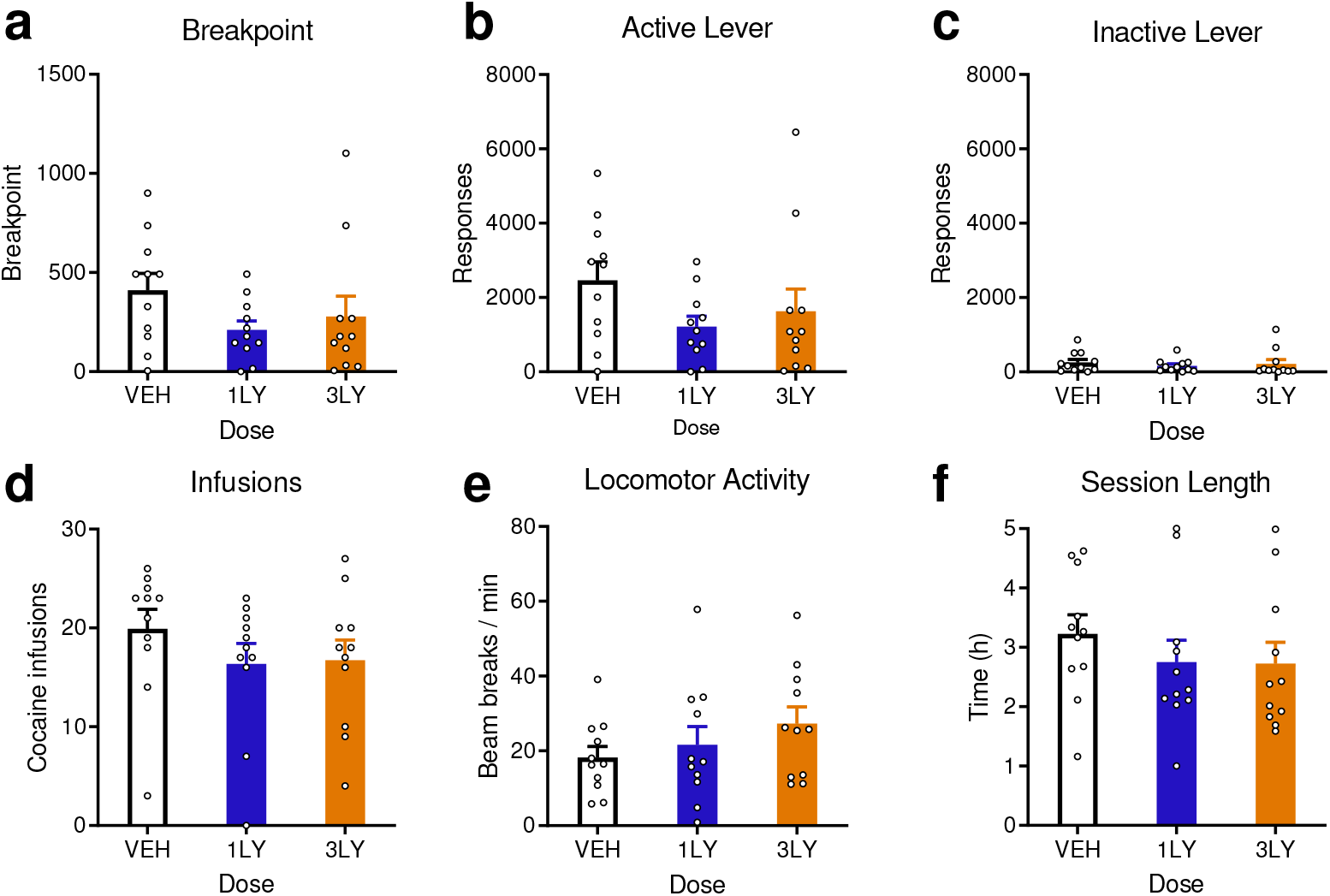
Effects of bilateral LY379268 microinjections into the dorsal striatum on responding for cocaine under a progressive ratio schedule of reinforcement. Following microinjection of 1 or 3 μg LY379268 into the dorsal striatum, rats (n = 11 males) showed a non-statistically significant tendency to reduce their (a) breakpoints for cocaine (p = 0.075) and (b) active lever presses (p = 0.058). (c) There were no significant effects of LY379268 microinjection on inactive lever presses. (d) LY379268 microinjection did not significantly change the number of cocaine infusions earned. There were no significant effects of LY379268 on (e) locomotion or (f) session length. Data are means ± SEM.

In summary, there was a statistically non-significant tendency for LY379268 to reduce breakpoints and active lever presses for cocaine under a PR schedule of reinforcement, in male rats. This, along with the presence of potential outliers at the highest LY379268 dose (See Figures 3a and 3c), warranted further study of the potential effects of the mGlu_2/3_ receptor agonist on responding for cocaine in a larger, mixed-sex cohort of rats.

### Experiment 2

#### Male and female rats self-administered similar amounts of cocaine during intermittent access sessions

After first reliably acquiring food and then cocaine self-administration behaviour, male (n = 11) and female (n = 10) rats now received ten, 5-h IntA cocaine self-administration sessions (Figure 4a). In contrast to Experiment 1, rates of lever pressing significantly changed across the 10 Sessions (F(9,171) = 2.16, *p* = 0.03). However, Bonferroni-corrected post-hoc comparisons did not indicate that any sessions were significantly different from any others (All *P*’s ≥ 0.39). There was also a significant Sex × Session interaction effect on lever-pressing behaviour (F(9,171) = 2.04, *p* = 0.038), but again Bonferroni-corrected post-hoc comparisons did not indicate that males and females differed on any specific sessions (*p*’s ≥ 0.24). All rats showed a strong overall preference for the active vs. inactive lever (F(1,19) = 54.76, *p* < 0.001). This preference was constant across session and biological sex, (no significant Lever × Session interaction effect; F(3.48,66.06) = 0.68, *p* = 0.58, ε = 0.39), Lever × Sex interaction (F(1,19) = 0.43, *p* = 0.52), or Lever × Session × Sex interaction (F(3.48,66.06) = 0.66, *p* = 0.6, ε = 0.39). There was also no significant main effect of Sex (F(1,19) = 0.16, *p* = 0.69), indicating that female and male rats showed similar levels of lever-pressing behaviour. Thus, across IntA sessions, rats reliably and consistently lever pressed for cocaine, and there were no significant differences between the sexes on this behavioural measure.

**Figure 4.**
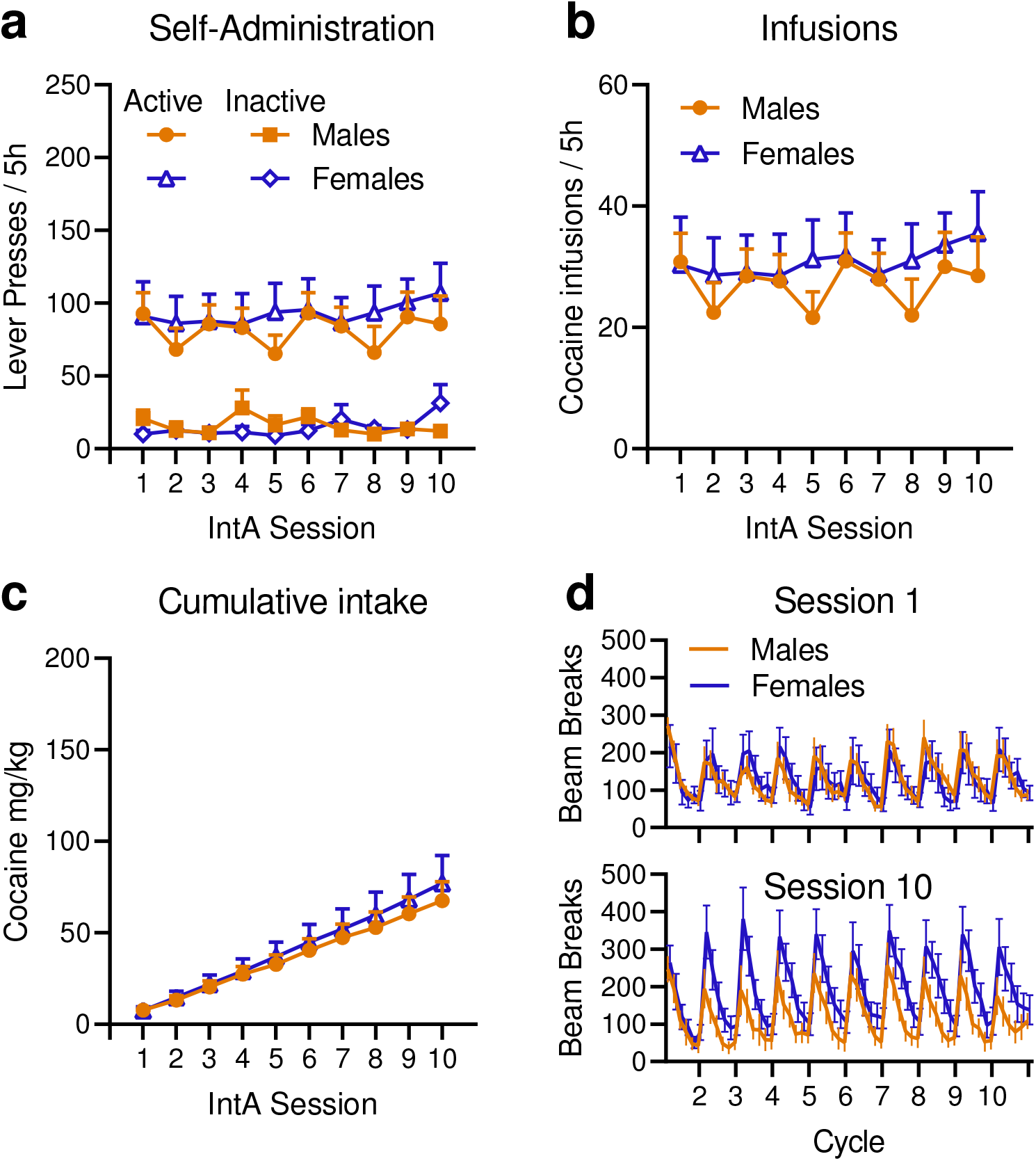
Intermittent access cocaine self-administration during Experiment 2. (a) Rats (n = 10 females; n = 11 males) showed stable lever pressing behaviour across the 10 IntA cocaine self-administration sessions, with a marked preference for the active vs. inactive lever. There were no significant differences between the sexes on these behaviours. (b) Rats earned a stable number of cocaine infusions across the IntA sessions, resulting in (c) a linear increase in cumulative cocaine intake across sessions. (d) During the 1st and 10th each IntA session, locomotor behaviour spikes at the beginning of each cocaine ON phase before decreasing during each cocaine OFF phase. Across the sexes, locomotion significantly increased across sessions, suggesting rats developed psychomotor sensitization. Data are shown as means ± SEM.

Female and male rats also consumed similar amounts of cocaine (Figure 4b; no main effect of Sex (F(1,19) = 0.27, *p* = 0.61) or Session × Sex interaction effect (F(3.88,73.74) = 0.96, *p* = 0.43, ε = 0.43). Consumption was stable across sessions (F(3.88,73.74) = 1.5, *p* = 0.21, ε = 0.43). This steady consumption resulted in a linear increase in cumulative cocaine intake (Figure 4c). Thus, female and male rats steadily self-administered cocaine across IntA sessions, with no significant differences between the sexes.

Female and male rats showed a similar locomotor response to self-administered cocaine across IntA sessions (Figure 4d; main effect of Sex F(1,19) = 0.52, *p* = 0.48); Sex × Session interaction effect (F(1,19) = 3.29, *p* = 0.099). Across the sexes, locomotor activity followed a spiking pattern, with peaks coinciding with cocaine ON phases, and locomotion decreasing during each cocaine OFF phase. In contrast to the smaller cohort of male rats tested in Experiment 1, the larger, mixed-sex cohort tested here showed evidence of psychomotor sensitization to self-administered cocaine. Mean locomotor peaks (in beam breaks / 5 min bin) were significantly higher in Session 10 than in Session 1 (Session: F(1,19) = 5.71, *p* = 0.027). Thus, while rates of cocaine intake remained stable across sessions, locomotor activity increased across sessions. This suggests that the rats developed psychomotor sensitization to self-administered cocaine, consistent with reports from our laboratory and others showing that IntA cocaine intake promotes the development of sensitization (Algallal et al., 2020; Allain et al., 2017; Allain & Samaha, 2018; Carr et al., 2020).

#### LY379268 reduced responding for cocaine under a progressive ratio schedule of reinforcement

Rats were first allowed to self-administer cocaine under a PR schedule, under baseline conditions (i.e., with no microinjections, indicated as ‘base’ or ‘baseline’ in Figure 5) and then following aCSF microinjections into the dorsal striatum (‘VEH’ or ‘Vehicle’). Compared to baseline, vehicle microinjection had no effect on breakpoints (Figure 5a; F(1,19) = 1.848, *p* = 0.19), lever pressing behaviour (Figure 5b; F(1,19) = 1.69, *p* = 0.209), self-administered infusions (Figure 5c; (F(1,19) = 0.027, *p* = 0.87) or session length (Figure 5d; F(1,19) = 0.52, *p* = 0.48). There were also no significant effects of Sex on these measures (Figure 5a; Sex F(1,19) = 0.54, *p* = 0.47; Treatment × Sex interaction effect (F(1,19) = 0.57, *p* = 0.46. Figure 5b; main effect of Sex F(1,19) = 0.8, *p* = 0.38, Treatment × Sex interaction effect F(1,19) = 1.52, *p* = 0.23, Lever × Sex interaction effect F(1,19) = 0.3, *p* = 0.59, Treatment × Lever × Sex interaction effect F(1,19) = 0.21, *p* = 0.65. Figure 5c; main effect of Sex F(1,19) = 0.27, *p* = 0.61, Treatment × Sex interaction effect F(1,19) = 0.32, *p* = 0.58. Figure 5d; main effect of Sex F(1,19) = 0.27, *p* = 0.61, Treatment × Sex interaction effect F(1,19) = 0.047, *p* = 0.83). Across biological sex and treatment condition, rats pressed more on the active vs. inactive lever (F(1,19) = 65.71, *p* < 0.001; Figure 5b). Finally, neither vehicle microinjection nor sex influenced locomotor activity during PR testing (data not shown; Treatment F(1,19) = 2.52, *p* = 0.13; Sex F(1,19) = 2.52, *p* = 0.13; Treatment × Sex interaction F(1,19) = 0.2, *p* = 0.66).

**Figure 5.**
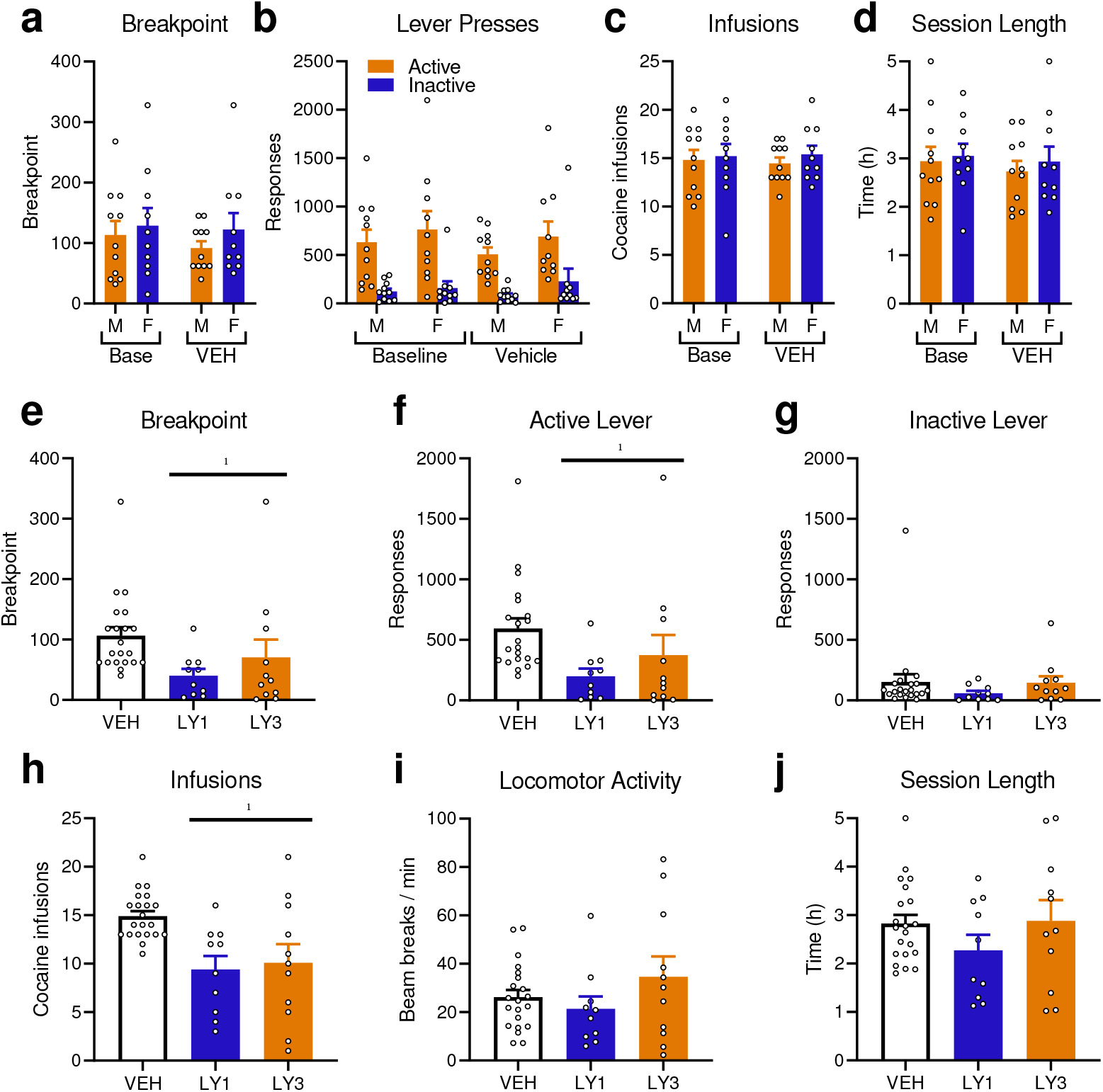
In female and male rats, LY379268 microinjections into the dorsal striatum significantly reduced responding for cocaine under a progressive ratio schedule of reinforcement. Relative to baseline conditions, where rats (n = 10 females; n = 11 males) received no microinjections, aCSF vehicle microinjection had no effects on (a) breakpoints for cocaine, (b) active or inactive lever presses, (c) cocaine infusions earned, or (d) session length. Compared to vehicle, LY379268 significantly reduced (e) breakpoint, (f) the number of active lever presses, and (g) cocaine infusions earned. LY379268 had no statistically significant effects on (h) inactive lever presses, (i) locomotor activity, or (j) session length. Data are means ± SEM.

Thus, relative to baseline, intra-dorsal striatum vehicle administration had no effect on responding for cocaine under a PR schedule of reinforcement. Female and male rats also showed similar responding for cocaine under PR conditions. As such, the sexes were pooled for subsequent analyses.

Relative to the vehicle test, LY379268 decreased multiple measures of responding for cocaine under PR conditions. LY379268 significantly reduced breakpoints (Figure 5e), active lever presses (Figure 5f), and self-administered cocaine infusions (Figure 5g). In each case, there was a main effect of Treatment, indicating a significant difference between vehicle and LY379268 sessions (Figure 5e: F(1,19) = 10.27, *p* = 0.005; Figure 5f: F(1,19) = 11.28, *p* = 0.003; Figure 5g: F(1,19) = 24.61, *p* < 0.001). The mGlu_2/3_ receptor agonist had similar effects on responding for cocaine at 1 and 3 μg/hemisphere (Figure 5e: main effect of Dose, F(1,19) = 0.21, *p* = 0.65, Treatment × Dose interaction effect (F(1,19) = 1.28, *p* = 0.27; Figure 5f: Dose effect, F(1,19) = 0.16, *p* = 0.69, Treatment × Dose interaction effect; F(1,19) = 1.49, *p* = 0.24; Figure 5g: Dose effect, F(1,19) < 0.001, *p* = 0.98, Treatment × Dose interaction effect; F(1,19) = 0.48, *p* = 0.5).These results suggest that under conditions where obtaining cocaine requires increasing effort, activation of dorsal striatal mGlu_2/3_ receptors reduces pursuit of the drug in male and female rats.

LY379268 had no significant effects on inactive lever presses (Figure 5h; no effect of Treatment F(1,19) = 1.42, *p* = 0.25, Dose F(1,19) = 0.83, *p* = 0.37, or Treatment × Dose interaction F(1,19) = 0.002, *p* = 0.96). Similarly, LY379268 also had no effect on locomotor activity relative to vehicle (Figure 5i; no effect of Treatment F(1,19) = 0.11, *p* = 0.74, Dose F(1,19) = 0.29, *p* = 0.59, or Treatment × Dose interaction F(1,19) = 3.73, *p* = 0.069). Mixed-design ANOVA revealed no significant differences in session length (Figure 5j) between vehicle and LY379268 tests, as there were no effects of Treatment (F(1,19) = 0.45, *p* = 0.51), no differences between 1 and 3 μg of LY379268 (Dose: F(1,19) = 2.04, *p* = 0.17), or Treatment × Dose interaction effect (F(1,19) = 0.28, *p* = 0.6). Thus, LY379268 left general motor function unchanged, suggesting that the decrease in instrumental responding for cocaine under a PR schedule of reinforcement involves suppression of the incentive motivational properties of cocaine.

## Discussion

We found that activating mGlu_2/3_ receptors in the dorsal striatum with LY379268 significantly reduced responding for cocaine under a PR schedule of reinforcement in female and male rats, without producing non-specific effects on motor function. In an initial experiment using male rats, we observed non-significant trends that were suggestive of a reduction in responding for cocaine during PR tests. In Experiment 2, which involved a larger, mixed-sex cohort of rats, intra-dorsal striatum LY379268 significantly reduced breakpoints achieved with cocaine reinforcement, total presses on the cocaine-associated lever, and cocaine infusions earned. Together, these results suggest that after a history of chronic cocaine use, activity at glutamatergic mGlu_2/3_ receptors in the dorsal striatum mediates incentive motivation for the drug.

### Effects of activating mGlu_2/3_ receptors in the dorsal striatum across experiments

There was variability in the response to cocaine between rat cohorts used in Experiments 1 & 2, but LY379268 had consistent effects on the pursuit of cocaine across experiments and also across male and female rats. Rats in Experiment 2 appear less sensitive to the rewarding effects of cocaine. During IntA sessions, they consumed approximately half as much cocaine as rats in Experiment 1 did (Figure 2b vs Figure 4b). Rats in Experiment 2 also achieved lower breakpoints for cocaine under a PR schedule (‘VEH’ conditions in Figure 3 vs Figure 5). Lower consumption might involve greater sensitivity of rats in Experiment 2 to the psychomotor activating effects of cocaine, as their peak locomotor behaviour was higher than that in rats in Experiment 1. Rats in Experiment 2— but not in Experiment 1— also showed evidence of locomotor sensitization across IntA sessions, with more locomotion on session 10 compared to session 1. These behavioural results suggest cohort differences in the sensitivity to cocaine. This could involve sourcing animals from different breeding colonies (Charles River Area C62 in Experiment 1 and R06 in Experiment 2), which can have significant effects on their behaviour (Khoo et al., 2022; Fitzpatrick et al., 2013; Gileta et al., 2022). Despite differences in the sensitivity to the rewarding and psychomotor effects of cocaine, LY379268 consistently suppressed the pursuit of cocaine across experiments (non-significant trends in Experiment 1 and significant suppression in Experiment 2). LY379268 also decreased responding for cocaine under a PR schedule in a mixed-sex cohort. Thus, the ability of LY379268 to reduce the pursuit of cocaine is robust against variability between cohorts of rats and is also seen across the sexes.

### Role of dorsal striatum mGlu_2/3_ receptors in incentive motivation for cocaine

Changes in glutamate homeostasis are thought to contribute to cocaine use disorder (Kalivas, 2009; Kalivas & Volkow, 2011; Spencer & Kalivas, 2017). mGlu_2/3_ receptors have an important role in glutamate homeostasis by providing negative feedback to glutamate neurons (Niswender & Conn, 2010; Linden et al., 2005) and can become dysregulated following cocaine use (Niedzielska-Andres et al., 2021). For example, cocaine self-administration increases mGlu_2/3_ expression in the dorsal striatum (Pomierny-Chamiolo et al., 2017; Beveridge et al., 2011). LY379268, which inhibits putative glutamate release (Kilbride et al., 1998; Dohovics et al., 2003), has also been shown to suppress PR responding for cocaine in rats (Allain et al., 2017; Hao et al., 2010). Prior studies have shown that mGlu_2/3_ receptors in the ventromedial prefrontal (or prelimbic) cortex (Shin et al., 2018) and in the nucleus accumbens core (Peters & Kalivas, 2006) mediate cue- and cocaine-induced cocaine seeking. To our knowledge, the present study is the first to implicate the mGlu_2/3_ receptors in the dorsal striatum. The current results therefore extend previous findings by identifying a role for mGlu_2/3_ receptor-mediated activity in the dorsal striatum in incentive motivation for cocaine.

### How LY379268 in the dorsal striatum might reduce incentive motivation for cocaine

We do not have direct evidence of LY379268’s mechanism of action. However, LY379268 is a highly selective and potent mGlu_2/3_ receptor agonist that can decrease presynaptic glutamate release (Conn & Pin, 1997; Imre, 2007; Schoepp, 2001; Monn et al., 1999; Farazifard & Wu, 2010). Thus, one possibility is that LY379268 reduces the pursuit of cocaine by attenuating glutamate release in the dorsal striatum. The dorsal striatum receives glutamatergic projections from several regions, including large glutamatergic inputs from cortical regions such as the orbitofrontal cortex as well as the mediodorsal and parafascicular nucleus of the thalamus (Hunnicutt et al., 2016; Peak et al., 2019; Wall et al., 2013). There are some reports that the mediodorsal thalamus is involved in cocaine-seeking behaviour (James et al., 2011; Weissenborn et al., 1998), and this could involve thalamic inputs to dorsal striatum. There is also evidence for a role of the orbitofrontal cortex and its projections to the dorsal striatum in the pursuit of cocaine. Pharmacological disconnection of the orbitofrontal cortex and dorsal striatum via contralateral microinjections of baclofen and muscimol suppresses responding for cocaine under PR conditions, suggesting reduced incentive motivation for the drug (Minogianis et al., 2019). It appears likely, given the findings of Minogianis et al. (2019) as well as other studies of the orbitofrontal cortex in appetitive motivation (Moorman, 2018; Cetin et al., 2004; Lucantonio et al., 2012; Stalnaker et al., 2015), that glutamatergic projections from the orbitofrontal cortex to the dorsal striatum are involved in mediating incentive motivation for cocaine. In addition to decreasing synaptic glutamate release, activation of mGlu_2/3_ receptors can also suppress excitability of projection neurons (Conn & Pin, 1997; Imre, 2007; Schoepp, 2001). As such, LY379268 could have reduced responding for cocaine by modulating neural activity in regions downstream of the dorsal striatum. Future studies using circuit manipulations can address the specific contributions of mGlu_2/3_ receptor-mediated effects in both pre- and post-synaptic dorsal striatum-dependent circuits.

## Conclusions

Microinjection of the mGlu_2/3_ receptor agonist LY379268 into the dorsal striatum significantly reduced incentive motivation for cocaine. In an initial, smaller experiment (Experiment 1), we observed non-significant reductions in responding for cocaine under a PR schedule of reinforcement. These effects were confirmed in a second, larger experiment using both male and female rats, where we observed a significant decrease in cocaine self-administration under PR conditions. Importantly, we did not observe any motor suppressive effects of LY379268, suggesting that the ability of increased dorsal striatum mGlu_2/3_ receptor activity to attenuate the pursuit of cocaine did not involve non-specific motor suppression. The effects of LY379268 were consistent across the sexes, and across cohorts that differed in sensitivity to the rewarding and psychomotor stimulating actions of cocaine. This suggests that the anti-addiction effects of LY379268 are robust, being evident in both females and males, and despite individual differences in the response to cocaine. Therefore, we suggest that signalling via mGlu_2/3_ receptors within the dorsal striatum mediates incentive motivation for cocaine.

